# BioTracs: A transversal framework for computational workflow standardization and traceability

**DOI:** 10.1101/2020.02.16.951624

**Authors:** Joséphine Abi-Ghanem, Djomangan Adama Ouattara

## Abstract

**Background:** The need of digital tools for integrative analysis is today important in most scientific areas. It leads to several community-driven initiatives to standardize the sharing of data and computational workflows. However, there exists no open agnostic framework to model and implement computation workflows, in particular in bioinformatics. It is therefore difficult for data scientists to share transparently and integrate heterogeneous analysis processes coming from different scientific domains, programing languages, projects or teams.

**Results:** We present here BioTracs, a transversal framework for computational workflow standardization and traceability. It is based on PRISM architecture (Process Resource Interfacing SysteM), an agnostic open architecture we introduce here to standardize the way processes and resources can be modelled and interfaced in computational workflows to ensure traceability, reproducibility and facilitate sharing. BioTracs is today implemented in MATLAB and available under open source license on GitHub. Several BioTracs-derived applications are also available online. They were successfully applied to large-scale metabolomics and clinical studies and demonstrated flexibility and robustness.

**Conclusions:** As an implementation of the PRISM architecture, BioTracs paved the way to an open framework in which bioinformatics could specify ad model workflows. PRISM architecture is designed to provide scalability and transparency from the code to the project level we less efforts. It could also be implemented using open object-oriented languages such as Python, C++ or java.

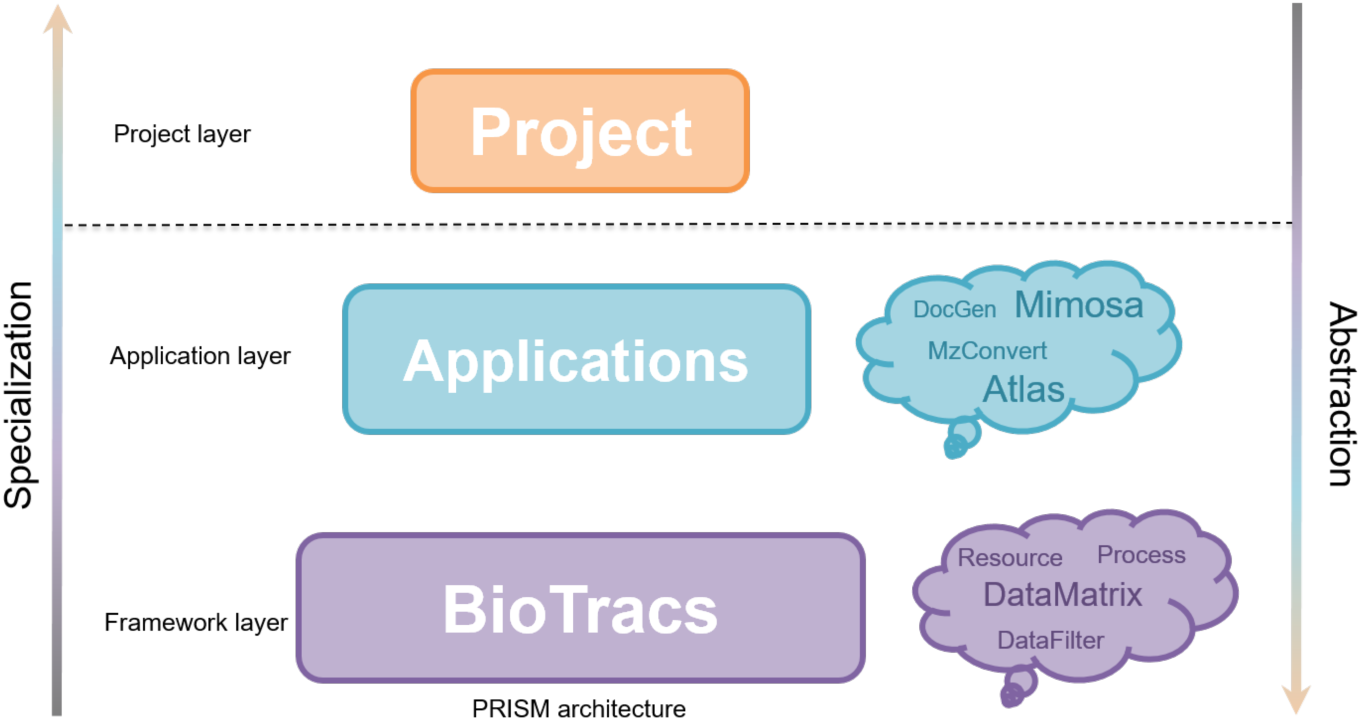

## Background

The need of digital tools for integrative analysis is today important in most scientific areas, in particular in bioinformatics where multi-source and heterogeneous data must be integrated in complex analysis processes for decision-making. Today, scientists from different scientific communities tend to merge their efforts to collaboratively solve challenging problems [1–4]. This trend is enhanced by the emergence of the Web 2.0 that facilitates the way people can share their ideas, experiences and scientific research all around the world. It particularly enhances the need of transparency (cross-verifiable workflows) to deliver quality results in research and development. The efficient use of these heterogeneous data and workflows relies on our capacity to structure them and standardize their annotation. This led to several initiatives for best practice convergences (data and workflow standardization) to allow experimental and computational scientists to easily perform, reproduce, and share their data and analyses. Standardization can be defined at different levels. The first one corresponds to resource standardization. Resources correspond to data used to feed analysis pipelines and generate industrial values. In non-computational areas, resources may refer to experimental samples used in laboratory experiments for instance. The second is process standardization. Processes correspond to algorithms or more complex experiments that are used to analyze resources and generate value. The third one is the standardization of projects’ annotation. Finally, standardization may also be defined at the level of application packaging to ensure that packaged processing engines function reliably and independently of the computing platform.

At the level of resource standardization, there exist several initiatives. The Gene ontology project (http://geneontology.org) aims at providing a framework for the annotation of gene functions and relationships. The HUPO proteomics standard initiatives (http://www.psidev.info/) aims at standardizing data annotation in metabolomics and proteomics [5–7]. The open source ISA framework aims at standardizing scientific resources, processes and projects [8–10] in several areas of life science. Web-based platforms such as Galaxy (http://galaxyproject.org/) or community-based computational initiatives such as Bioconductor (http://www.bioconductor.org), Biopython (https://biopython.org/) rather focus on the standardization the sharing of processes, in particular how computational scientists can package and distribute computing toolboxes in bioinformatics [2, 11, 12]. Some developments oriented toward clinical data integration and sharing such as tranSMART and eTriks are also concerned (http://transmartfoundation.org/, http://www.etriks.org/) [3]. More recently, one assisted to the emergence of containerization systems to ensure the reliability of the packaging and sharing of computational applications. They rely on docking systems (e.g. Docker, Virtual environments) that work like virtual machines to ensure that each docked application (a container) can be executed safely and generate reproducible results in different environments. For instance, the Anaconda (https://www.anaconda.com) and BioContainer (https://biocontainers.pro/) projects are community-driven projects that provide guidelines and repositories to create and distribute containers in data science and bioinformatics, respectively [4]. All these initiatives are driven by a need of reproducibility, transparency and traceability in computational sciences.

However, it remains difficult for data scientists to transparently share and integrate heterogeneous analysis codes coming from different scientific domains, projects or teams. Generally, the use of a given toolbox written in a given language (R, Matlab, Python, etc.) requires dealing with non-standard interfaces. For example, different Galaxy instances can rapidly diverge and become incompatible although they are based on the same initial platform. ToolSheds were therefore introduced to solve such divergences during the implementation of *ad hoc* Galaxy tools on different Galaxy instances. They serve as “appstores” to all Galaxy instances worldwide. The Bioconductor project is probably among those that provide a rigorous computational framework for sharing R programs in bioinformatics [11]. Its success is based on open and clear rules for submitting R codes with standard annotations (i.e. well-documented codes).

Despite these improvements, it can be difficult to interface different building blocks coming from different teams, in particular if different data formats are used. In these conditions, custom data parsers and exporter may be required to adapt the building blocks. It therefore lacks today an agnostic (i.e. non-language specific) framework to help computational scientists to standardize, annotate and share their computational processes at the code level. To address this issue, we first introduce here PRISM (Process Resource Interfacing SysteM), an agnostic open architecture that standardizes the way processes and resources can be interfaced in computational workflows, to ensure traceability, reproducibility and facilitate sharing. This framework is not only devoted to computational experiments, but can be extended to model any experimental process. It is based on a high-level abstraction to provide an agnostic and transversal framework that allows integrating processes and resources coming from heterogeneous communities while allowing computational scientists to derive custom workflows that fit to their specific requirements. Contrary to existing frameworks, it is specifically designed to fully capture real-world processes while ensuring simplicity at the code level. We therefore present BioTracs, which is an implementation of the PRISM architecture. BioTracs is a computational framework implemented in MATLAB (The MathWorks) and it delivers a set of libraries to implement building blocks in bioinformatics. It also provides libraries to wrap and use language-specific programs without compromising the integrity of the whole system. BioTracs provides ready to used templates classes to created complex applications. BioTracs library is freely available online on GitHub (https://bioaster.github.io/biotracs/).

## Implementation

### Core architecture: fundamental concepts

Only two basic concepts are necessary for modeling computational workflows. The first one is the *process* that contains the algorithmic logic. The second one is the concept of *resource* that represents input data and output data (i.e. results) used and generated by processes, respectively. PRISM is only based on these two concepts and proposes an agnostic architecture to interface processes and resources in complex computational workflows (Figure 1). A process is moreover associated with a *configuration* to store analysis parameters. One configuration instance can be associated with several process instances. Inputs and outputs are interfaces composed of a set of ports. A port is allowed to contain only one resource. To ensure traceability, a resource must refer to its creator process and this one must exist. This design therefore introduces a strong coupling between a resource and its creator and has several consequences in terms of transparency. In particular, one of the rules of the PRISM architecture is that only processes are allowed to create or alter resources. To improve flexibility and code reusability we introduce here the notion of *engine*. An engine is a black-box process that can be internally used by a process to achieve some subtasks. As it is shown later, engines are not used to build workflows, but are meant to improve code reusability and task delegation without caring about the traceability of the subtasks that are delegated. In the PRISM architecture, a process may therefore be composed of zero to several engines as presented by Figure 1.

**Figure 1:**
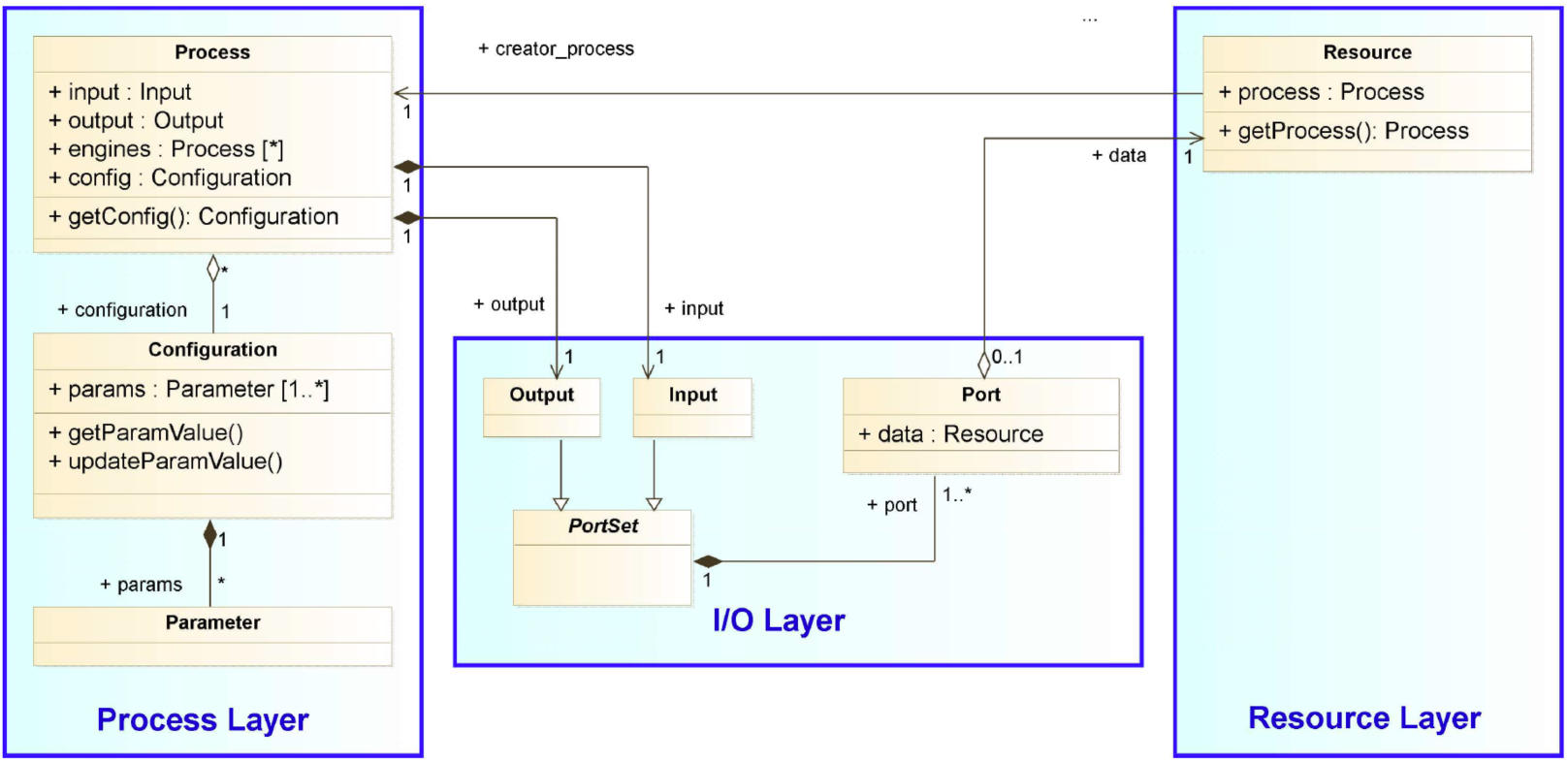
PRISM architecture. PRISM relies on a process layer and resource layer that are linked through an input/output layer. Input and output objects inherit from Interface and thus contain a data (resource).

In practice, the implementation of PRISM must guarantee the safe execution of the processes, to store and visualize the resources that are used and produced at each steps of a workflow. PRISM was therefore built on the basis of the well-known MVC (model-view-controller) architecture. In the MVC architecture, a model contains information (e.g. metadata) that are displayed or stored in a database. A view is responsible for managing human-machine interfaces and interactions (e.g. the visualization of plots, tables, graphs, button clicks, …). The controller implements the *actions* that are execute when the user interacts with the model and the view (e.g. an action to launch a workflow, to display a plot). This architecture therefore separates the role and responsibilities of these three components in the system (Figure 2). In PRISM, the *Process*, the *Configuration* and the *Resource* inherit from the base *Model* class because these objects store user and system data. For example, a *Process* stores system metadata associated with the state and the execution of an algorithm. A *Configuration* stores user parameter values. *Resource* stores the analysis data and result values. These objects can therefore be associated with several views. For example, a dataset can be visualized as a table, a series of plots, a heatmap or a graph depending on the context. Basically, models and views are generally sufficient to implement simple and complex workflows. In PRISM, the controller was introduced to model high-level business logics when implementing user actions in applications as described in the BioTracs-MIMOSA application (manuscript submitted).

**Figure 2:**
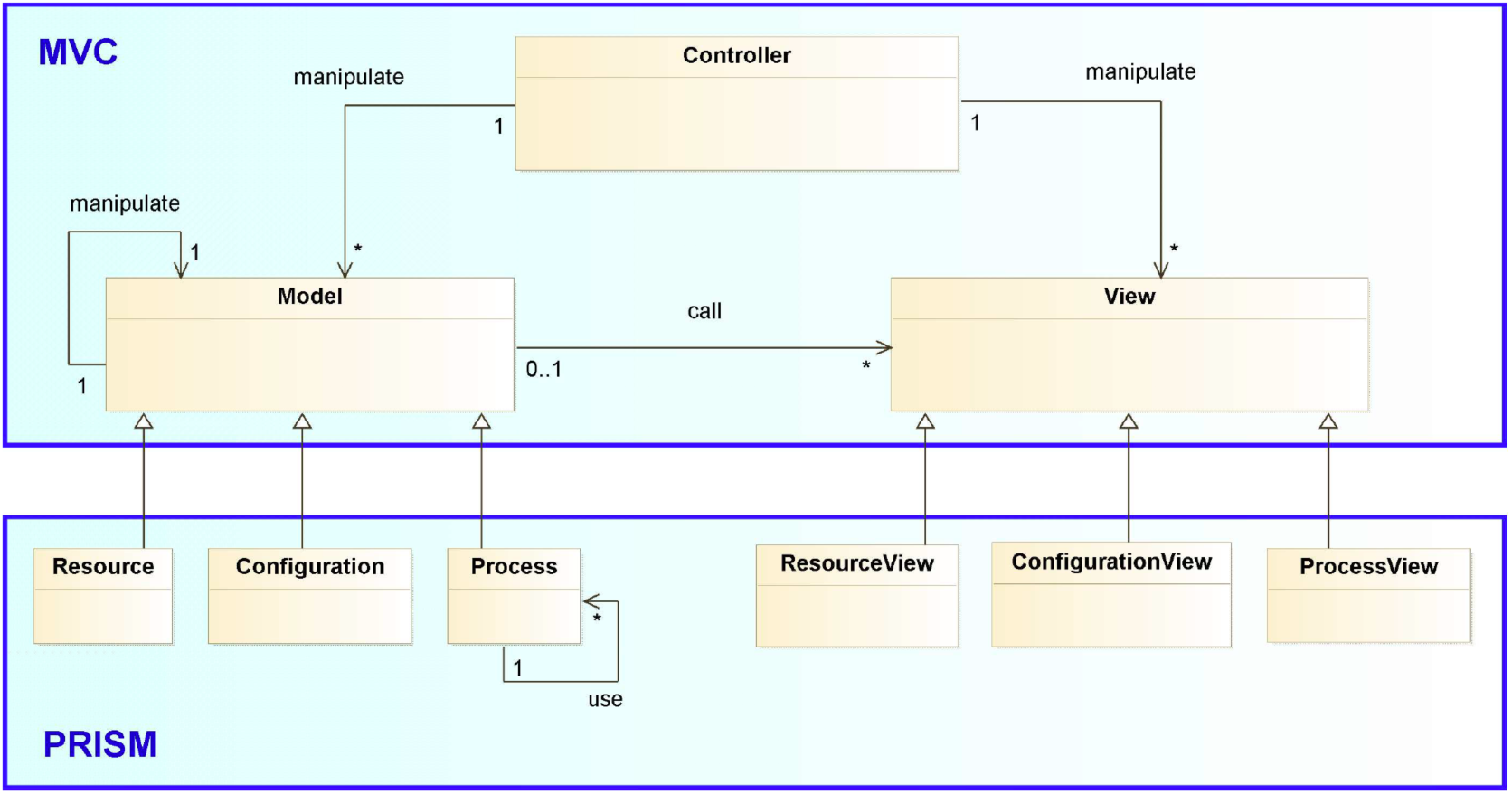
PRISM components are implemented on top of an MVC architecture. Resource, Configuration and Process inherit from Model and are associated with ResourceView, ConfigurationView and ProcessView, respectively.

A workflow is a process chain in which processes are connected through their input and output ports (Figure 3). It is a process containing sub-processes. It is then modelled using a hierarchical structure (Figure 4). The Workflow class therefore inherits from the Process class and is composed of a set of Process nodes.

**Figure 3:**
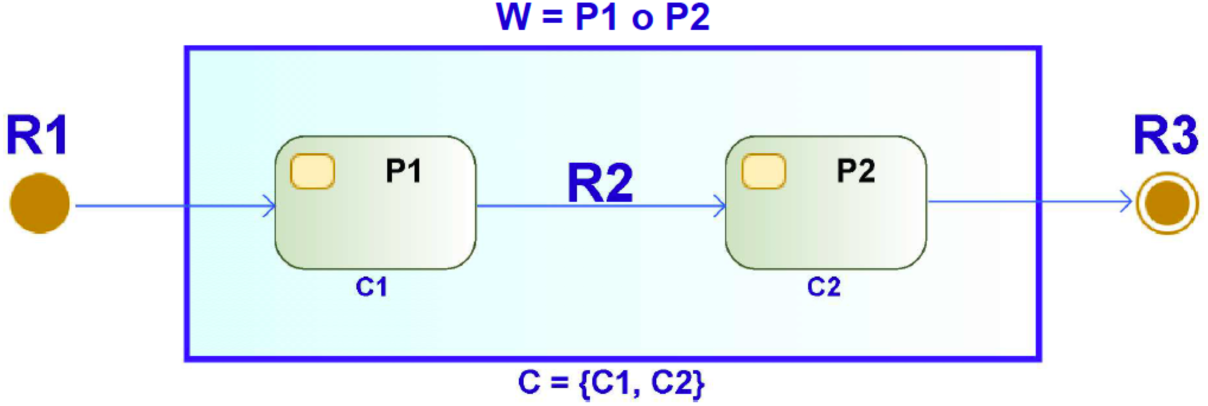
Illustration of a workflow composed of two processes. Process P2 is left-compatible with F1 or equivalently process P1 is right-compatible with P2. C1 and C2 are configurations associated with processes F1 and F2 respectively. C is the configuration of the workflow. R1, R2 and R3 are resources.

**Figure 4:**
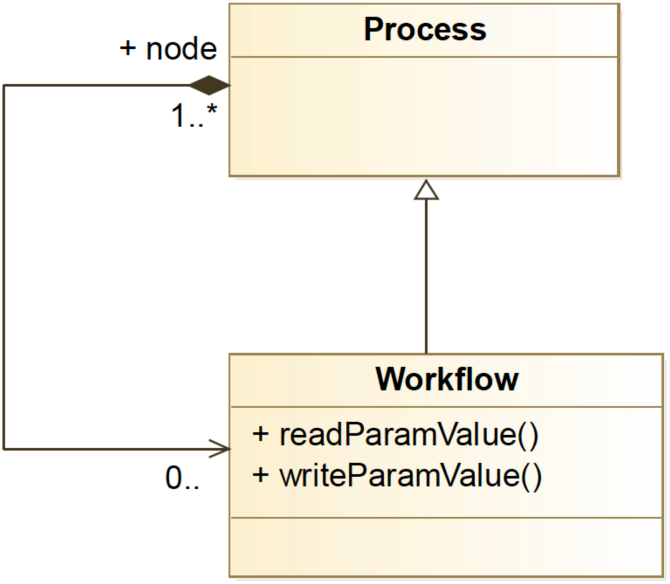
A workflow is as process that contains a set of process nodes. A least one process is needed to build a workflow. Methods readParamValue and writeParamValue are used to calibrate the workflow.

Compatible nodes can be chained in the workflow through their input and output ports. In practice, the configuration of workflows is challenging, in particular when the user wants to access and configure all sub-processes in a hierarchical manner. The Workflow class provides interfaces methods (readParamValue and writeParamValue methods) to directly configure sub-processes (Figure 4).

### BioTracs framework implementation

PRISM is an abstract concept and is usable in practice only if it is implemented. BioTracs is an implementation of the PRISM architecture in MATLAB and is based on a strong experience acquired in computational metabolomics. It defines several building blocks that inherit PRISM thinking. The BioTracs framework provides components to implement bioinformatics workflows. These building blocks are useful to create custom applications such as BioTracs-Mimosa in metabolomics, BioTracs-Polaris in proteomics and BioTracs-Atlas for biostatical analyses. The BioTracs framework is available on GitHub under an open source license (https://bioaster.github.io/biotracs/).

## Results & Discussions

Several analytic platforms and standardization initiative were created to standardize the way computational scientist work together. These initiatives are fueled by the emergence of big data and powerful solutions based on artificial intelligence that leverage these data to generate new insights. We therefore propose here the BioTracs framework that is an implementation of the PRISM architecture, which is introduced here. The main advantage of the PRISM architecture is its agnosticism and its simplicity because it well captures real-world processes. The BioTracs framework was used to create, in a scalable manner, metabolomics applications that were successfully used in academic and industrial projects involving large-scale clinical metabolomics data [13]. The BioTracs-Mimosa application is one of them and is presented in a paper submitted in BMC bioinformatics. BioTracs-Mimosa provides ready-to-use workflows managed by a controller that a user can access and configure the workflows. After execution, the user can inspect workflows and their sub-processes (logs, internal configurations, results and result views).

### A 3-layer architecture scalable from the code to the project level

System scalability is the ability to extend a system to new purposes. It relies on well-designed architectures in which the components can be safely upgraded without compromising the whole system. That requires that components are loosely coupled with well-defined interfaces. Loose coupling reduces dependencies between components, i.e. each component must have a limited knowledge about the internal functioning the other ones. PRISM defines how resources are interfaced with processes. Any upgrade of a given node (process) in a workflow does not require upgrading the other processes as long as the adjacent nodes remain compatible. In addition, an essential property required to ensure system scalability is abstraction. Abstraction consists in identifying components that share common functions and factorizing these functions in more general (more abstract) components (Figure 5). For instance, in the library of BioTracs-Spectra (https://github.com/bioaster/biotracs-m-spectra), the NMR and MS spectra were modelled using the same mother class *DataMatrix* because they can both be represented as N-by-2 matrices, where N is the number of m/z and ppm ticks for MS and NMR, respectively. The two columns represent the m/z (or ppm) axis and the intensity axis (see classes *NMRSpectrum* and *MSSpectrum*). Processes that are compatible with the *DataMatrix* class will be compatible with the *NMRSpectrum* and *MSSpectrum* classes by inheritance. Abstraction is therefore a powerful mean for code reusability. The main advantage of this architecture is to bring functionalities from the code to the business level, while ensuring the role and responsibility of all the stakeholders in the development process (e.g. developers, non-coder experts, clients). BioTracs is three-layer system in which core BioTracs libraries are specialized to create applications such as BioTracs-Spectra, BioTracs-Mimosa, BioTracs-Atlas for instance.

**Figure 5:**
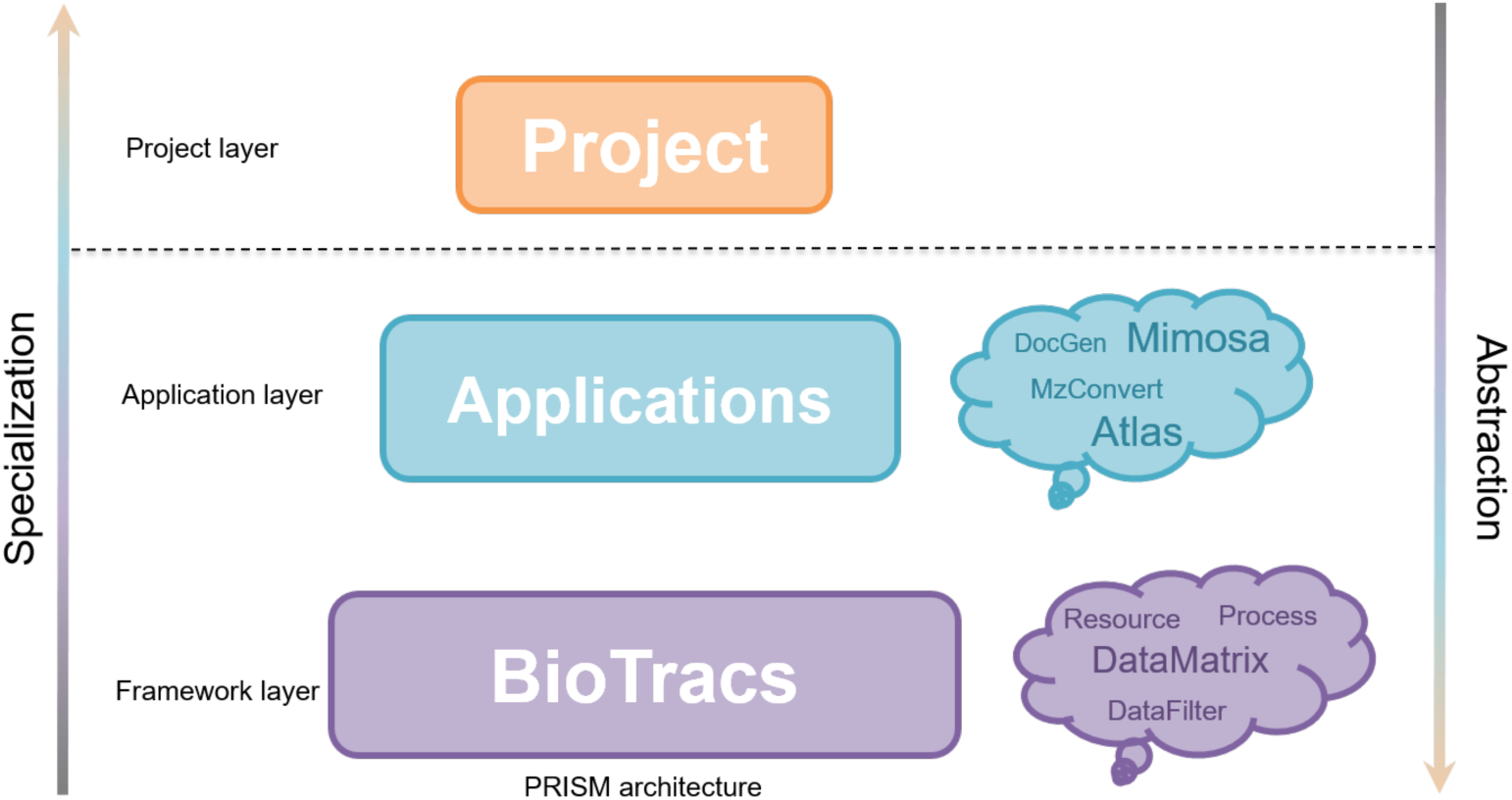
A 3-layer system scalable from the code to the project level. BioTracs is based on the PRISM architecture. Application layer inherits from and specializes core BioTracs libraries. The Project layer contextualizes and uses applications.

### Comparison with existing standardization systems

Several solutions were developed in the past to improve the sharing and the standardization of the computational tools. These standardization systems apply at different levels: at the level of the management of workflows (KNIME, Snakemake, Airflow, etc.), the packaging of applications (Docking systems, R Bioconductor), the standardization of the input/output data (file formats standardization), the standardization of the description of experiments (e.g. ISA tabs frameworks). The R Bioconductor framework is based on a package submission process to promote high-quality, well documented, and interoperable software in bioinformatics. For instance, R Bioconductor packaging promotes the use of common R data structures (S4 classes and methods) to easily operate with existing infrastructure. One on the most recent approach that is gaining interest in bioinformatics is the use of docking systems to build containers (a.k.a. biocontainers). Containers allow to package applications in order to simplify their deployment and use without worrying about dependencies in heterogeneous platforms. This platform, supported by the ELIXIR consortium (https://biocontainers.pro), is not language specific and ensures analysis reproducibility. It does provide not guideline on the implementation of computational processes. PRISM is not devoted to replace docking systems and BioTracs processes could be “dockerized” to create containers to facilitate their sharing. Compared to R Bioconductor, BioTracs provides a more rigorous implementation architecture not only focused on data structure but also processes and workflows.

Workflow management systems such as KNIME and Snakemake allow encapsulating and chaining programs as building blocks to easily author, monitor and schedule workflows. They offer advanced management systems for cloud computing and analysis parallelization on clusters. BioTracs is, by construction, a system that allows chaining MATLAB algorithms and even external programs (R, Pyhton, Java) to build complex workflows. However, PRISM was not designed to be an advanced workflow management system. It does not define how to distribute and optimize the parallelization of workflows on cluster nodes. It is rather designed to standardize earlier, in the development process, the implementation of computational building blocks and provide a universal scheme to design complex workflows. It therefore allows to easily contribute to the development of computational applications independently of the way these applications could be packaged, deployed or executed (e.g. using docking systems or more complex workflow management systems). It is therefore well adapted to multi-disciplinary environments were different scientists could be federated in the development and maintenance of the same computational pipelines. PRISM shares some similarities with the ISA tab framework [9, 14]. In the ISA framework, an *Investigation* file is used to describe the overall goals and means used in an experiment or a project. It is a high-level concept used to link several studies and corresponds to the business layer. *Studies* are components that describe *subjects* under study (e.g. patients, biological samples characteristics, protocols and parameters). This layer therefore encapsulates to the notion of *Resource* in PRISM (in particular biological resources), but also the associated protocols applied to generate these resources. *Assays* describe tests performed on *subject materials* in order to produce data (e.g. use of a mass-spectrometer a acquisition system to generated data). They therefore correspond to notion of *Process*, but at the level of the acquisition instruments (Figure 6). The functioning of PRISM-based frameworks such as BioTracs can therefore be modelled using the ISA framework. BioTracs framework is therefore complementary to the ISA framework.

**Figure 6:**
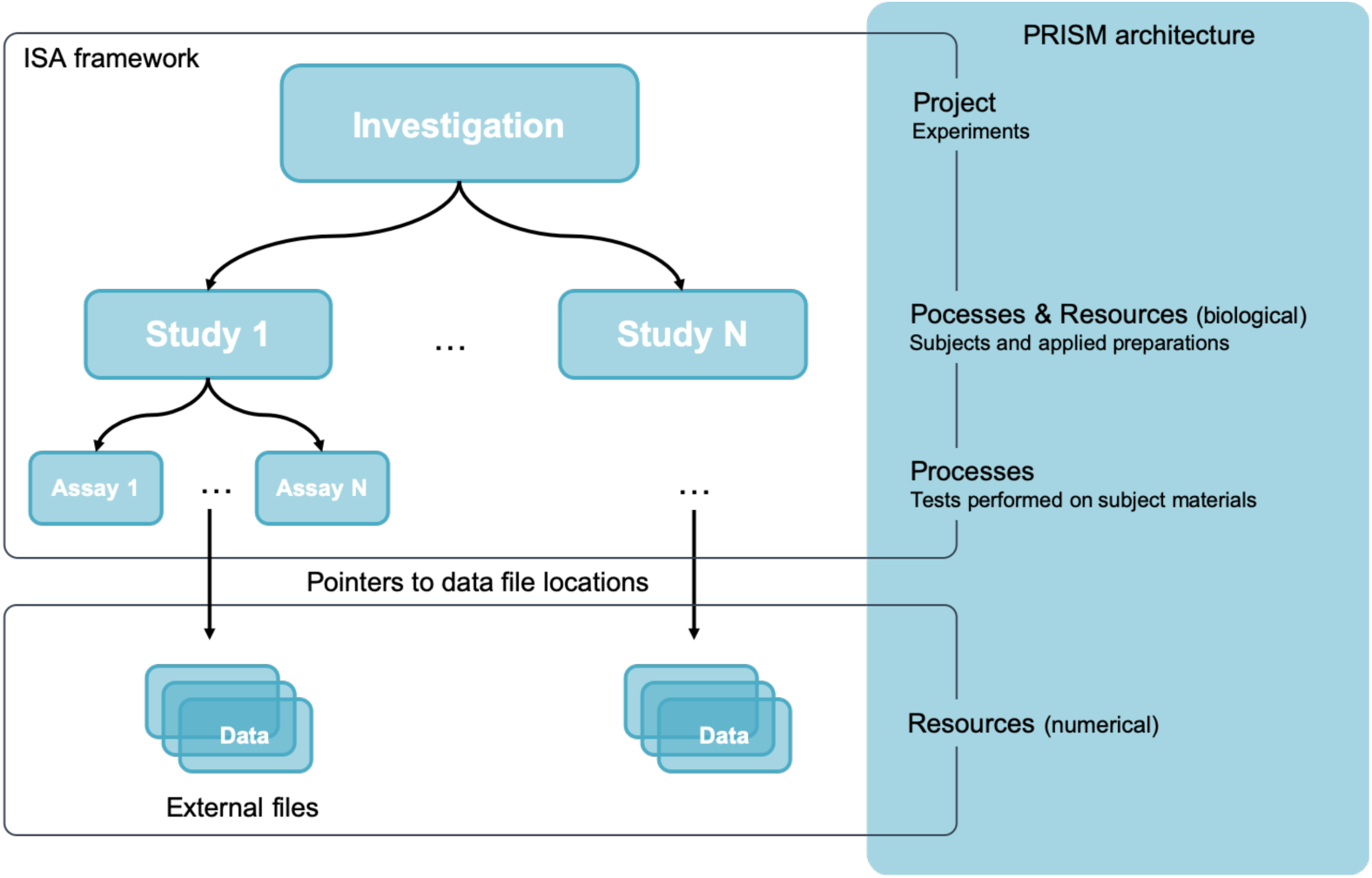
The ISA framework and its links with the PRISM architecture.

### Application of BioTracs to a toy analysis workflow (TAW)

We illustrate here, through a simple toy analysis workflow (TAW), how BioTracs can be used. Consider an initial dataset where rows and columns are observations and variables respectively. Each observation belongs to a group. The user is interested in performing a basic unsupervised analysis on this dataset to extract trends in the data, followed by a supervised analysis to extract features that discriminate each observation group. Let us build a simple analysis workflow to achieve this task (Figure 7). This workflow uses some basic statistical modules implemented in the core BioTracs and in the BioTracs-Atlas application devoted to machine learning. The first process is a *FileImporter* (FI) used to import a disk file as an in-memory resource file (*DataFileSet*) corresponding to the dataset. The *DataParser* next parses the resource file to create the *DataSet* object. The *Filter* is used to remove variables having negligible variances (i.e. that can be considered constant) or having all their values under a predefined limit of quantitation. It namely prevents having ill-conditioned problems in the following numerical solvers. The *DiffProcess* performs differential analysis using Student t-test. The *PCALearner* process performs unsupervised analysis using principal component analysis. The *PLSLearner* process performs classification using partial least square discriminant analysis (PLS-DA) [15, 16]. The *PLSPredictor* process receives the *PLSLearnerResult* and the initial training dataset as input resources to perform prediction. Also, a blind test dataset could be used to assess the learned model on a new set of observations. The *PartialDiffProcess* process performs “partial differential analysis” in the principal components space computed from the PCA. It is based on the Mahalanobis distance that is suitable to measure group separation in multivariate dimensional spaces [17, 18]. *PartialDiffProcess* also computes a Fisher-test based p-value to assess the significance of group separations. Partial differential analysis therefore generalizes univariate t-test to multivariate problems. In BioTracs-Atlas, the *PCALearner* and *PLSLearner* derived from a mother class called *BaseDecompLearner* because they belong to the class of eigenvalue decomposition problems. The *PartialDiffProcess* can therefore assess group separations in the PCA or PLS score plots. Next, the user visualizes the analysis results using custom views generated and exported by the *ViewExporter* (VE) as images (.jpg, .png, etc.) of MATALB figures (.mat). *FileExporter* (FE) processes allow writing in-memory resources as disk files than can next be imported for further analyses. For traceability purposes, each process, as well as the encapsulating workflow, generate individual configuration files (*config.params.xml*) containing all the parameters used at each stage of the workflow as well as log files. TAW code file is available online on GitHub (https://github.com/bioaster/biotracs-papers). It provides three datasets for testing: the well-known Fisher’s Iris flower dataset [19], the breast cancer Wisconsin dataset [20, 21] and an anonymized “metabo” dataset we constructed for classification analysis testing. The metabo dataset consists in 30 samples belonging to five groups (A, B, C, D and E). Each sample is characterized by a set of 46 metabolic features generated using NMR. Figure 8 presents the views generated by the main steps of the workflows using the metabo dataset. Views and results obtained with the three datasets are available online on GitHub.

**Figure 7:**
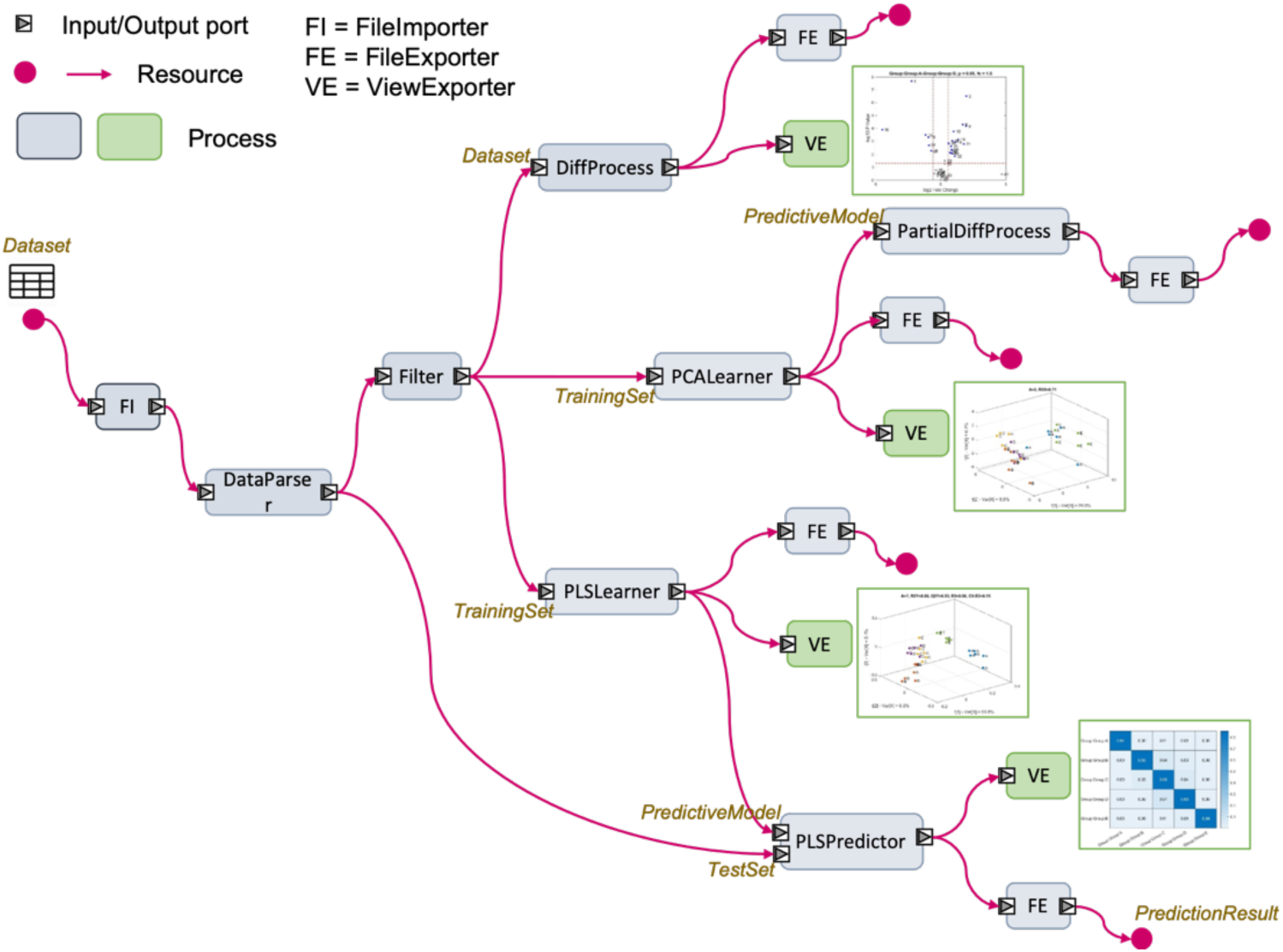
Graphical illustration of TAW (toy analysis workflow). Resources are sketched using bullets and arrows (resource flow).

**Figure 8:**
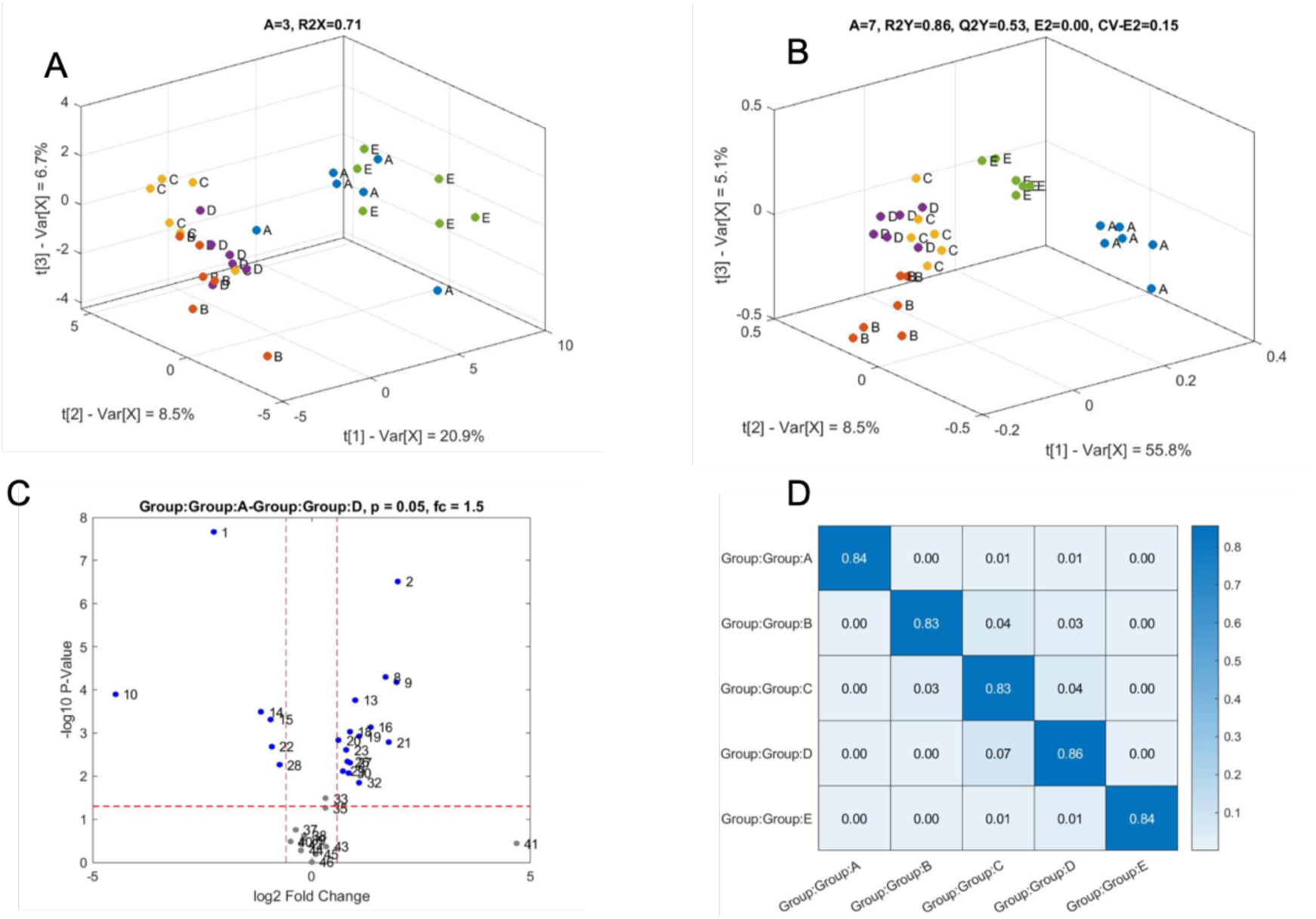
Views exported by the Toy Analysis Workflow using the metabo dataset. (A) PCA 3D score plot. (B) PLS 3D score plot. (C) Volcano plot generated after the differential analysis showing most significant features when comparing group A vs group D. (D) confusion matrix generated after the PLS predictor process.

## Conclusions

We presented BioTracs, a transversal framework for the standardization and traceability of computational workflows. Fundamental concepts underlying BioTracs were presented. BioTracs derives from the PRISM architecture, which allows modeling and interfacing computational resources and processes to build complex workflows. It is designed to provide scalability and transparency from the code to the project level and demonstrated flexibility and robustness in large-scale metabolomics and clinical projects [22]. It provides a framework in which computational scientists can implement simple building-block processes and next specialize them to fit their scientific use-cases. Today, BioTracs and related libraries are implemented in the object-oriented MATLAB environment. It can be extended to open object-oriented environments such as Python, C++ or java.

## Availability and requirements

**Project name:** BioTracs

**Project home page:** https://bioaster.github.io/biotracs/

**Operating system(s):** Windows

**Programming language:** Matlab

**Other requirements:** None

**License:** BIOASTER Open Source

**Any restrictions to use by non-academics:** Please contact BIOASTER

## List of abbreviations

Not applicable

## Declarations

### Ethics approval and consent to participate

Not applicable

### Consent for publication

Not applicable

### Availability of data and materials

The BioTrac framework is available on GitHub. Please visit https://bioaster.github.io/biotracs/

The datasets and code used in the current study are available in the GitHub repository, https://github.com/bioaster/biotracs-papers in subfolder “demo”.

### Authors’ contributions

ADO wrote the paper. ADO and JAG wrote the code. ADO supervised the work.

## Acknowledgements

The authors would to acknowledge Frédéric BEQUET for its advices during the development of BioTracs.

